# CTCF-mediated Chromatin Structures Dictate the Spatio-temporal Propagation of Replication Foci

**DOI:** 10.1101/525915

**Authors:** Qian Peter Su, Ziqing Winston Zhao, Luming Meng, Miao Ding, Weiwei Zhang, Yongzheng Li, Mengzhu Liu, Rongqin Li, Yi-Qin Gao, Xiaoliang Sunney Xie, Yujie Sun

**Affiliations:** State Key Laboratory of Membrane Biology, Biomedical Pioneering Innovation Center (BIOPIC), School of Life Sciences, Peking University, Beijing 100871, China.; Graduate Program in Biophysics and Department of Chemistry and Chemical Biology, Harvard University, Cambridge, MA 02138, U.S.A.; Institute of Theoretical and Computational Chemistry, College of Chemistry and Molecular Engineering, Peking University, Beijing 100871, China.

## Abstract

Mammalian DNA replication is initiated at numerous replication origins, which are clustered into thousands of replication domains (RDs) across the genome. However, it remains unclear whether the replication origins within each RD are activated stochastically. To understand how replication is regulated at the sub-RD level, we directly visualized the spatio-temporal organization, morphology, and *in situ* epigenetic signatures of individual replication foci (RFi) across S-phase using super-resolution stochastic optical reconstruction microscopy (STORM). Importantly, we revealed a hierarchical radial pattern of RFi propagation that reverses its directionality from early to late S-phase, and is diminished upon caffeine treatment or CTCF knockdown. Together with simulation and bioinformatic analyses, our findings point to a ‘CTCF-organized REplication Propagation’ (CoREP) model. The CoREP model suggests a non-random selection mechanism for replication activation mediated by CTCF at the sub-RD level, as well as the critical involvement of local chromatin environment in regulating replication in space and time.

---

The integrity of genomic information is maintained across generations through DNA replication, a fundamental process that occurs in every mitotic cell during S-phase of each cell cycle. After decades of effort in biochemistry and genetics, we now have a comprehensive understanding of the molecular mechanisms of DNA replication as well as the components of the replication machinery^1,2^. Moreover, we also possess substantial insight into the genomic organization of DNA replication in the sequence space, in which replication is initiated at discrete loci along the chromosomes known as replication origins, which demarcate the genome into multiple replicons^3,4^. Such organization is manifested in physical space as thousands of spatially clustered sites or puncta within the nucleoplasm, termed replication foci (RFi) or ‘factories’, as revealed by fluorescence imaging^4–6^. Recent advances in high-throughput sequencing techniques have also enabled studies of DNA replication on a genome-wide scale. One such major breakthrough is the discovery of replication timing program, which revealed that different regions of the genome tend to replicate in a temporally specific manner^7–9^. Such temporally distinct replication regions, termed replication domains (RDs), could be considered as the counterpart of RFi in sequence space^6^. In fact, cells respond and adapt to cellular environment by modulating the timing of RDs for specific genes, but the mechanisms remain elusive^10,11^.

Several previous studies have suggested that the initiation of DNA replication is regulated at the RD level^9,12,13^, while within a RD, the activation is rather flexible and stochastic^14,15^. However, direct measurements on spread-out DNA fibers by DNA combing experiments have shown that a RD harbors several replicons that fire synchronously^10,16^, thereby implicating a possible activation mechanism dictated by the spatial organization of chromatin. Such proposal is also in line with the fact that the genomic boundaries of RDs align well with those of topologically associating domains (TADs)^17^, a prevalent structural feature of chromatin^18^.

In light of these findings, a fuller understanding of replication activation within the RD requires *in situ* monitoring and profiling of DNA replication sites in the context of the surrounding chromatin microenvironment at the sub-TAD level, with single-cell sensitivity and high spatiotemporal resolutions. Among the technical tools available to address this problem, sequencing-based methods are mostly limited to analyzing a population of cells, and thus only suitable for studying DNA replication at the RD level. On the other hand, majority of previous imaging studies are limited in their spatial resolution as well as their characterization of related functions, and are thus unable to provide details into the regulation of replication activation and propagation within RD. More recently, super-resolution microscopy has been utilized as a powerful tool for studying nucleosome heterogeneity^19^ as well as the morphological features of chromatin DNA in different epigenetic states^20–22^.

In this study, we use stochastic optical reconstruction microscopy (STORM) to probe the spatio-temporal organization of DNA replication within individual RFi at sub-diffraction-limit resolution in single human cells. Using metabolic labeling, we quantitatively characterized the morphology, sequence length, and replicon number of individual RFi across different stages of S-phase, lending support to the ‘increasing efficiency model’ of replication origin activation. We also characterized, for the first time, the *in situ* epigenetic signatures of individual RFi across S-phase, and demonstrate the correlation between replication timing and chromatin modification and organization at the RD level. Importantly, we observed an intriguing spatio-temporal propagation pattern within each replication focus, with DNA that are replicated earlier spatially surrounded by DNA that are replicated later during early S-phase. The radial directionality of such propagation pattern is reversed in late S-phase, and diminished upon treatment with caffeine (an inhibitor of checkpoint kinase) or knockdown of CTCF, a key organizer of chromatin architecture. Together with simulations and bioinformatic analysis, our results point to a ‘CTCF-organized REplication Propagation’ (CoREP) model, in which DNA replication activates non-randomly within individual RFi, mediated by CTCF-organized chromatin structures. These findings shed critical new insights into the regulation of origin activation at the sub-RD level, and have broad implications for the spatio-temporal coordination of DNA replication across the genome, which may be applicable to a variety of mammalian systems.

## RESULTS

### Super-resolution imaging and quantitative characterization of RFi across S-phase

To label newly replicated DNA *in situ*, we first synchronized cells to the G1/S boundary as previously described^23,24^. At specific stages of S-phase, a short 30-min pulse of dye-labeled dUTP or EdU was then supplied to the cells (see Methods for details), which incorporated these analogue molecules into the DNA synthesized during the pulse periods^25^. Imaging the labeled RFi with STORM (**Fig. S1a**) revealed a punctate distribution as well as changing physical morphology across S-phase (**Fig. 1a**), similar to previous observations using immuno-gold electron microscopy^26^, conventional optical microscopy^16,27^ and different super-resolution microscopy^21,28–31^. However, the superior spatial resolution afforded by STORM (~20 nm) enabled us to perform more accurate quantitative characterization of RFi, particularly for findings based on multi-color colocalization analysis.

**Figure 1.**
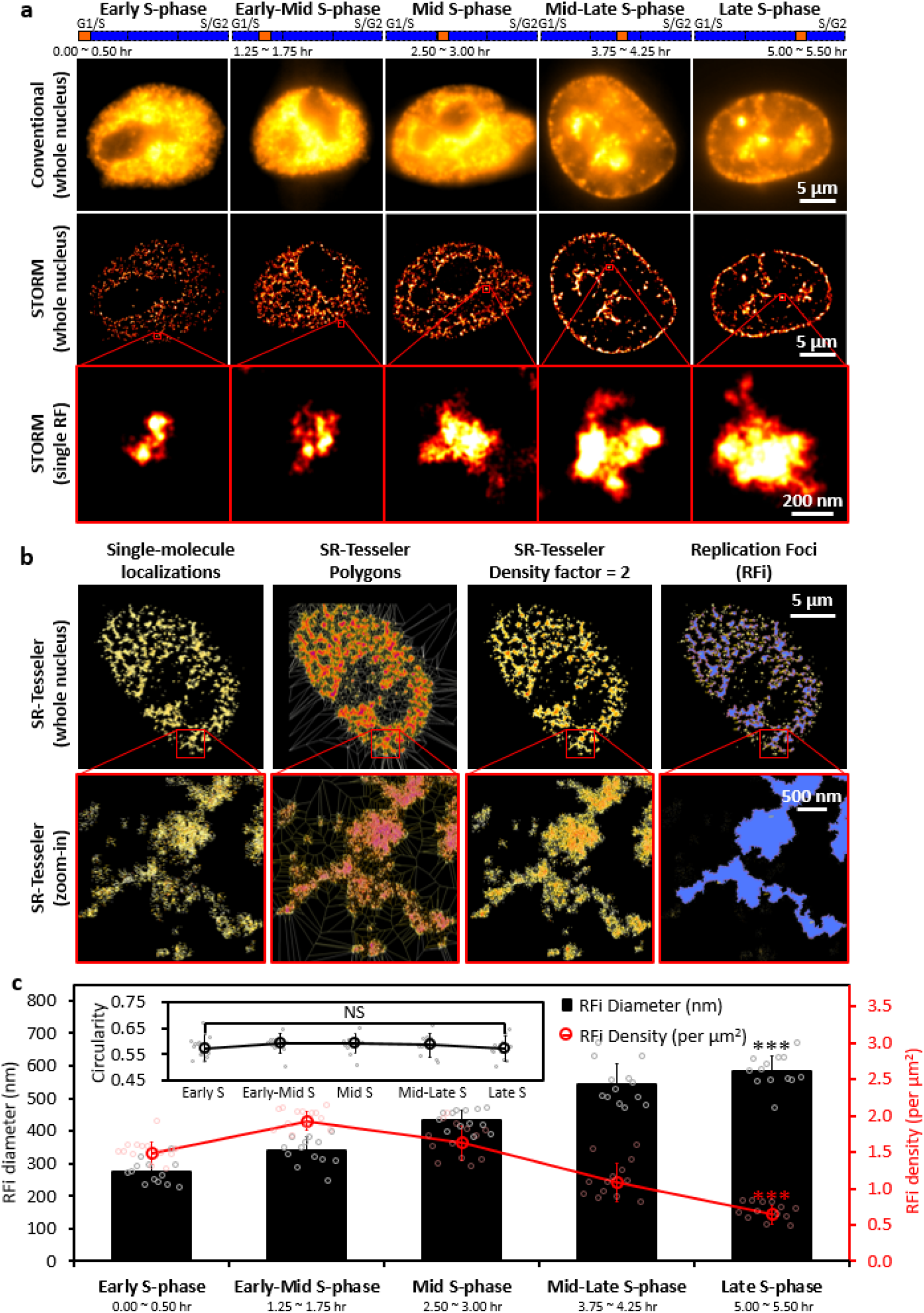
STORM imaging and quantitative characterization of replication foci (RFi) across S-phase. **a**, Conventional and STORM images of RFi in single HeLa cell nuclei at five different stages (from left to right column, each with 30min of labeling) across S-phase. Insets show zoomed-in representative images of individual RFi (in red boxes) at each stage. The orange boxes at the top indicate the relative temporal position of the labeling periods at respective stages of S-phase (blue bars, not drawn to scale). **b**, Unbiased hierarchical identification of RFi with SR-Tesseler, starting from raw single-molecule localizations in STORM images, to polygons in SR-Tesseler, to polygons after using density factor of 2, and finally to RFi. Insets show zoomed-in areas (in red boxes). **c**, Diameter (black column), density (red line), and circularity (inset) of RFi at each of the five stages imaged in **(a)**. Error bars denote mean ± s.d. p values are determined by Student’s t test; NS: not significant; ***: p < 0.001. (n = 10~13 cells for (c)). Source data are provided as a Source Data file.

To do so in a robust and unbiased fashion, we adopted a recently developed segmentation strategy, SR-Tesseler^32^, to group the detected single-molecule localizations into RFi (**Fig. 1b**). While RFi density peaks during early-mid S-phase and slowly declines during late S-phase (**Fig. 1c**, red curve), RFi size increases steadily from a mean value of 275.7 ± 35.6 nm during early S-phase to 582.7 ± 48.7 nm during late S-phase (**Fig. 1c**, black column). In contrast, the circularity of the foci, defined as the ratio between the semi-minor and semi-major axes of the foci, remains relatively unchanged from early to late S-phase (**Fig. 1c**, inset). We also ascertained that our synchronization and labeling procedures minimally impacted the growth kinetics of the cells as well as the morphology of the RFi (**Fig. S1b** and **c**).

As a validation for our imaging approach, we correlated the RFi observed in Cartesian space with the actual replicated DNA in sequence space using DNA combing approach^33^ (**Fig. 2a**). By imaging stretched single DNA fibers *in vitro*, we found that each single-molecule localization in the STORM images corresponded to 51.6 ± 12.6 base pairs of replicated DNA (**Fig. 2b**). This calibration allowed us to quantify the sequence length of RFi *in situ* for the first time (see Methods). Interestingly, we found that the mean sequence length of RFi produced during the 30-min pulse labeling increases from a mean value of 177.8 ± 19.6 kb in early S-phase to 950.02 ± 70.03 kb in late S-phase (**Fig. 2c**). These results are in agreement with previous sequencing measurements of 400 to 800 kb per replication domains (RDs)^17^ and 880 kb per topologically associating domains (TADs)^18^, lending support to the validity of our observations.

**Figure 2.**
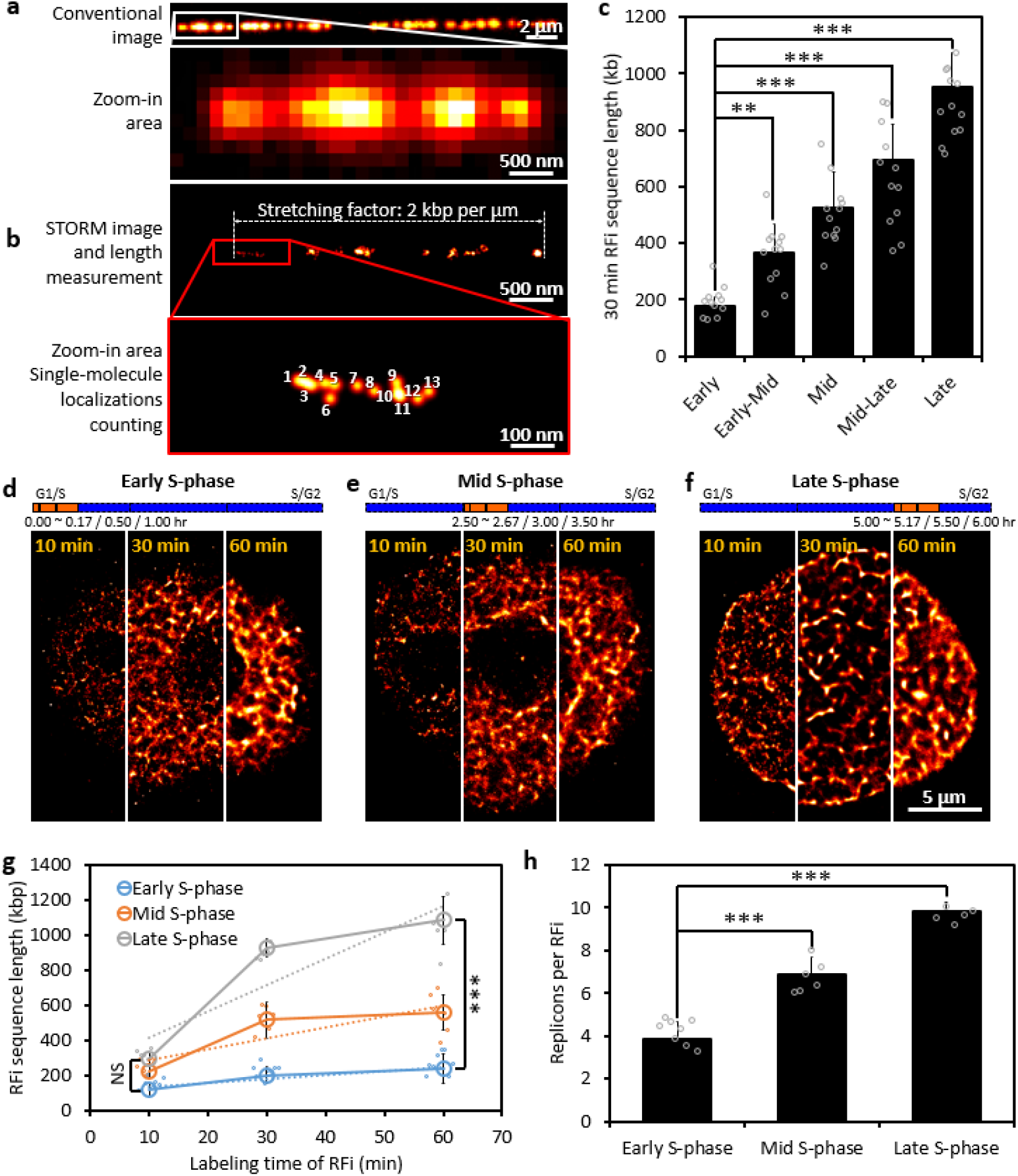
Sequence length and replicon number of individual RFi during different stages of S-phase. **a, b**, Conventional **(a)** and STORM **(b)** image and their zoomed-in areas of Alexa 647-EdU-labeled replication sites on a single stretched DNA molecule. The physical length of the DNA molecule can be directly measured and converted into base pair length^29,37^. Insets show zoomed-in areas; The singlemolecule localizations in the STORM images can be directly counted. c, The sequence length associated with individual RFi during each of the five stages imaged in **Fig. 1a. d-f**, STORM images of cells with their newly replicated DNA labeled for 10 min, 30 min or 60 min respectively from the beginning of early **(d)**, mid **(e)**, and late **(f)** S-phase. Each of the three images is stitched from 3 cells labeled for 10 min, 30 min and 60 min. The orange boxes at the top indicate the relative temporal position of the labeling periods at respective stages of S-phase (blue bars, not drawn to scale). **g**, Plot of the sequence length of RFi for different labeling duration at early (blue), mid (orange) and late (gray) S-phase and their linear fits (dashed lines). **h**, The in situ number of replicons per replication focus from early to late S-phase. Error bars denote mean ± s.d. p values are determined by Student’s t test; NS: not significant, ***: p < 0.001. (n =10~12 cells for (c) and 5~8 cells for **(g)** and **(h)**). Source data are provided as a Source Data file.

Moreover, by varying the pulse labeling duration (from 10 min, 30 min to 60 min) during early, mid, and late S-phase (totally 9 cell samples), we also monitored the rate of expansion of the size and sequence length of individual RFi (**Fig. 2d** to **f** for STORM images from 9 cells, **Fig. S1a** for the RFi identified, **Fig. 2g** for RFi sequence length distribution, and **Fig. S2b** for the RFi area distribution), and estimated the replication speed (defined as sequence length per replication focus per minute, a linear fit of the plot in **Fig. 2g**). While the area and sequence length of RFi are similar for short labeling duration (10 min), they increase much faster during late S-phase for longer labeling durations (30 and 60 min). Using the replication fork speeds (defined as sequence length per fork per minute) obtained previously from DNA combing experiments^34^, we further found that 3.84 – 9.82 replicons exist inside each RF at different stages of S-phase (**Fig. 2h**), and the origin efficiency increases from early to late S-phase, thereby supporting the ‘increasing efficiency model’ of DNA replication^35,36^.

### RFi across S-phase are associated with distinct stage-specific epigenetic signatures *in situ*

Even though the relationship between DNA replication timing and their chromatin environment has been studied by sequencing and biochemistry approaches, the measurements are often performed on cells in their G1-phase, lacking *in situ* spatial and dynamic information in S-phase^10,22,38^. To investigate the dynamic changes of epigenetic surrounding during replication propagation at sub-diffraction-limit resolution, we co-labeled RFi and seven key nuclear markers associated with a variety of chromatin signatures *in situ* using two spectrally distinct dyes, Alexa 647 and Cy3B, and performed two-color STORM imaging to quantify their colocalization (**Fig. 3a to g**, in which the contrast between the two colors has been enhanced for better visualization, see also **Fig. S3a** and **b** for conventional and normal contrast SR images, **Fig. S3c** and **d** for zoomed-in areas of caryoplasm and nuclear periphery). Among these markers, CTCF is known to regulate 3D chromatin architecture by forming chromatin loops and defines the boundaries between euchromatic and heterochromatic DNA^39–41^. Histone modifications H3K27ac and H3K4me3 are associated with transcriptionally active chromatin, with the former being a marker for active enhancers and the latter for active promoters^42,43^. In contrast, nuclear lamina (NL)-localized lamin A/C is suggested to mostly associate with transcriptionally repressed DNA^44^, while SUZ12 constitutes part of the polycomb repressive complex 2 (PRC2)/EED-EZH2 complex, which methylates Lys-9 (H3K9me) and Lys-27 (H3K27me) of histone H3, leading to transcriptional repression of the affected target genes^45,46^. Lastly, proliferating cell nuclear antigen (PCNA) is an essential component of the mammalian replication fork and acts both as a processivity factor for DNA polymerase δ and as a recruiter of other proteins involved in DNA replication^47^. Thus, it serves as a positive control together with H2B, which provides an indication of the general histone background of the nucleus.

**Figure 3.**
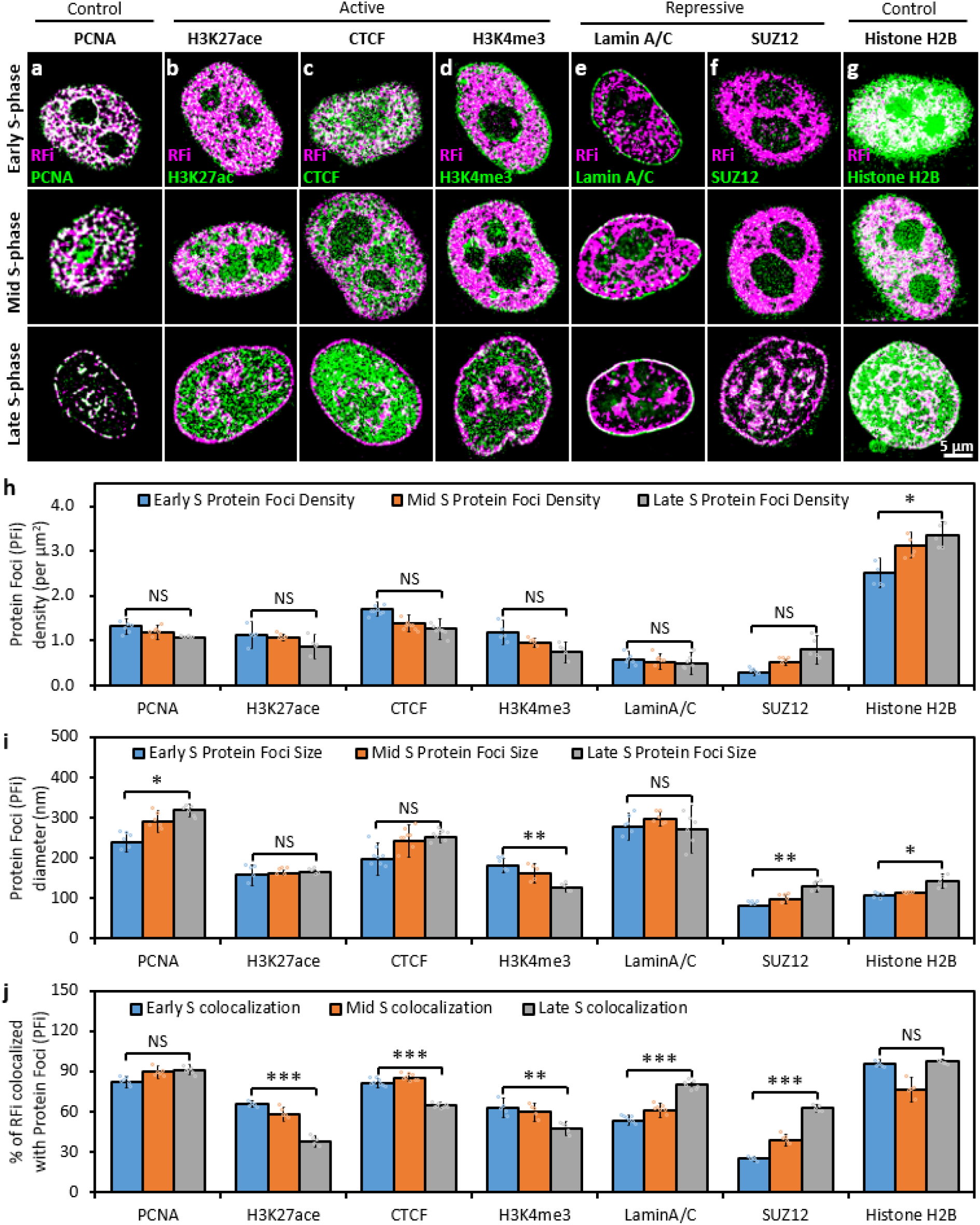
Dynamics of RFi-associated epigenetic signatures in situ across S-phase. **a-g**, Colocalization analysis between RFi and seven key nuclear markers reveals distinct chromatin states of RFi across S-phase. Cells with newly replicated DNA (pink) labeled at the beginning of early (top row), mid (middle row), and late (bottom row) S-phase were subsequently immunolabeled with antibody against each marker (green), and imaged with dual-color STORM. Contrast between the two colors was enhanced here for better visualization. See also **Fig. S3** for corresponding images with normal contrast as well as zoomed-in areas from each image. **h, i**, Density **(h)** and size **(i)** of protein foci (PFi) associated with each of the seven markers imaged in **(a-g). j**, Fraction of RFi that colocalize with the PFi of each marker during early (blue column), mid (orange column), and late (gray column) S-phase, respectively. Error bars denote mean ± s.d. p values are determined by Student’s t test; *: p < 0.05; **: p < 0.01; ***: p < 0.001; NS: not significant. (n = 5~8 cells for **h** to **j**). Source data are provided as a Source Data file.

Quantitative analysis of STORM images revealed that, while the densities and sizes of the protein foci (PFi) do not vary significantly across S-phase (**Fig. 3h** and **i**), the extent of RFi colocalized with each marker changes drastically (**Fig. 3j**, see also **Fig S3e** and **f** for a heat map and comparison to a conventional image). The relatively constant high-level colocalization ratio of RFi with PCNA (**Fig. 3a**) was indicative of the essential role of PCNA in DNA replication throughout S-phase, along with histone H2B (**Fig. 3g**), lending support to the accuracy of our colocalization analysis. Specifically, the fractions of RFi that colocalized with H3K27ac (**Fig. 3b**), CTCF (**Fig. 3c**), or H3K4me3 (**Fig. 3d**), all of which associate with the more open and transcriptionally active chromatin, showed a marked decrease from early S-phase to late S-phase (**Fig. 3b** to **d**). In contrast, the fractions of RFi that colocalized with lamin A/C (**Fig. 3e**) or SUZ12 (**Fig. 3f**), which are involved in lamina-associating domains (LADs)^48^ and transcriptional repression, displayed a pronounced increase from early to late S-phase (**Fig. 3e** and **f**). These results indicate that the DNA sequence near open and transcriptionally active chromatin tend to be replicated during early in S-phase, while those near closed and transcriptionally inactive chromatin tend to be replicated during mid and late S-phase. Collectively, our imaging results constitute the first *in situ* characterization of the dynamic epigenetic signatures of RFi, providing important evidence for the correlation between DNA replication timing and chromatin modification.

### The spatio-temporal dynamics of RFi propagation are directionally specific

To probe the spatial organization and dynamics of individual RFi, we next set out to investigate the dynamic propagation patterns of DNA replication in space and time. To do so, newly replicated DNA during two consecutive 30-min time windows at three stages of S-phase were labeled with two spectrally distinct dyes, Alexa 647 and Atto 550. See Methods for labeling details and the comparison of these two dyes in **Fig. S4a**. In addition, lamin A/C in the nuclear lamina was immunofluorescently labeled with a third dye, Atto 488 (**Fig. 4a** to **c**). Upon multi-color STORM imaging of the labeled cells, the relative spatial distributions of the foci during both time windows revealed a unique pattern of RFi spatial progression. During early S-phase, DNA synthesized during the second window occupied a greater area and enveloped DNA synthesized during the first window (**Fig. 4a**, top row of inset); this trend could be better visualized by enhancing the contrast between the two colors (**Fig. 4a**, bottom row of inset). 3D STORM imaging excluded the possibility that the colocalization pattern observed in 2D images resulted from objects that did not overlap in the z-direction (**Fig. S4b**). Significantly, this spatial propagation pattern became less pronounced during mid S-phase (**Fig. 4b**), and reversed its radial directionality in late S-phase (**Fig. 4c**), during which the DNA synthesized in the first window enveloped the DNA synthesized in the second window and occupied a greater area.

**Figure 4.**
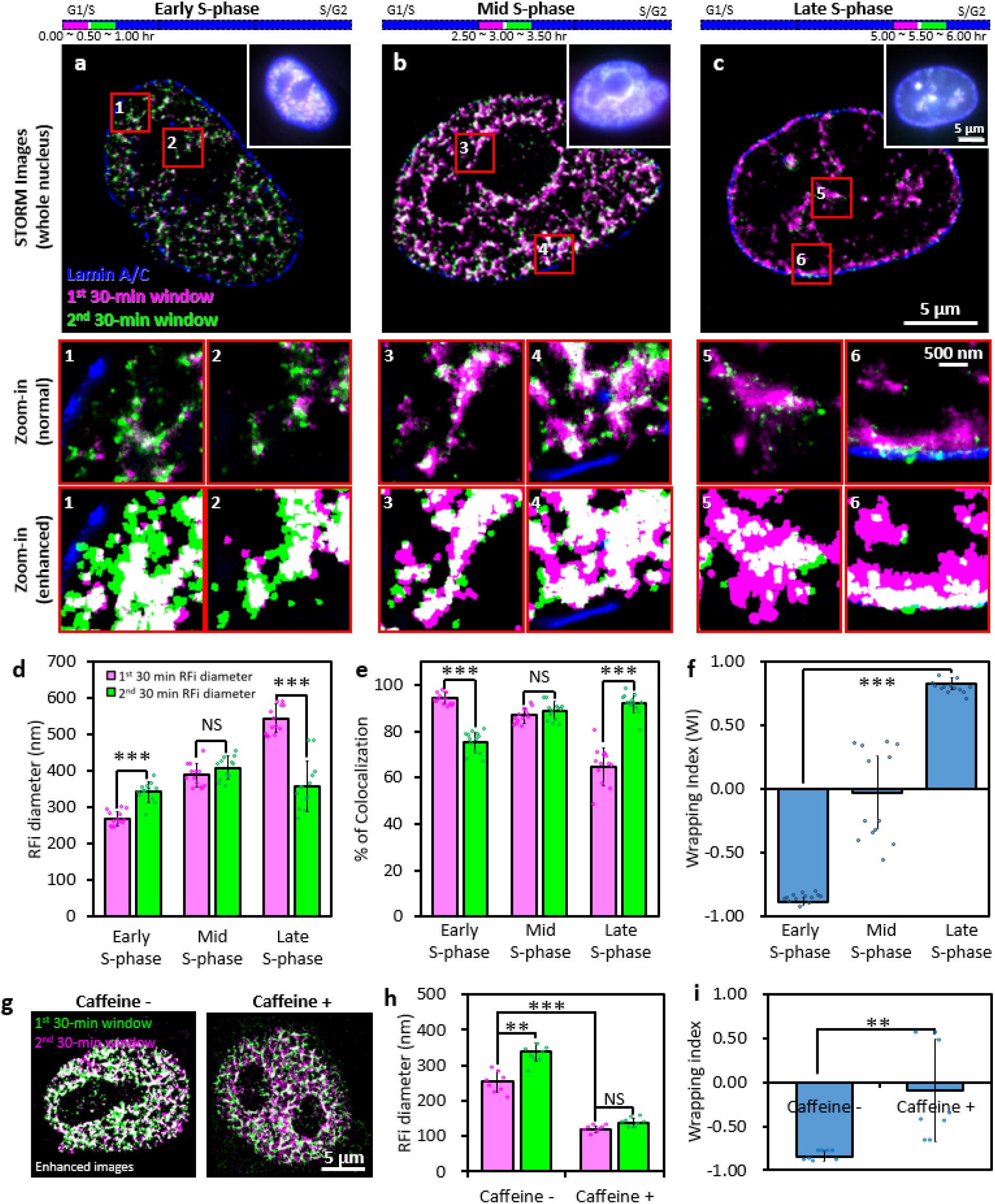
RFi during early and late S-phase display opposing spatio-temporal patterns of propagation dynamics. **a-c**, Newly replicated DNA during two consecutive 30-min windows at the beginning of early **(a**, 0~1.0 hr), mid **(b**, 2.5~3.5 hr), and late **(c**, 5.0~6.0 hr) S-phase were labeled (purple for the first 30-min and green for the second 30-min, white means the colocalized RFi), together with lamin A/C (blue) to demarcate the nuclear boundary of each cell. Multi-color STORM images of single nuclei are shown in comparison with conventional images (top right corner inset) of the same nuclei, and blue bars at the top indicate the relative temporal position (orange bars) of the labeling windows during the respective stages of S-phase (not drawn to scale). Insets (numbered 1 through 6) show zoomed-in areas (in red boxes), with the middle row showing normal contrast and the bottom row showing enhanced contrast between the two colors for better visualization. **d**, Diameters of RFi produced during the first 30-min (purple) and second 30-min (green) labeling windows at early, mid, and late S-phase. **e**, Fractions of RFi in the first labeling window that colocalize with those in the second window (pink), and vice versa (green), at each of the three stages of S-phase. **f**, Wrapping Index (WI) between the RFi replicated during the two labeling windows at each of the three stages of S-phase. **g**, Enhanced images of newly replicated DNA in two consecutive 30 min windows (purple and green represent for 1^st^ and 2^nd^ 30 min labeling, respectively) at the beginning of early S-phase with or without 5 mM Caffeine treatment. **h**, The presence of caffeine, which is well-known to inhibit the checkpoint and slow down the replication fork, results in the decrease of RFi diameter. **i**, Wrapping Index (WI) between the RFi replicated during the two labeling windows with or without caffeine treatment at early S-phase. Error bars denote mean ± s.d. p values are determined by Student’s t test; ***: p < 0.001; **: p < 0.01; NS: not significant. (n = 13 cells for **(d)** to **(f)**, and n=8 for **(h)** to **(i)**). Source data are provided as a Source Data file.

Such propagation dynamics can be more quantitatively illustrated by measuring the diameters of RFi in both time windows (**Fig. 4d**) as well as the extent of colocalization between them (**Fig. 4e**). Alternatively, to better quantify the radial directionality of RFi spatial propagation, we defined a wrapping index (WI) as 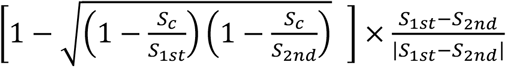, where *S*_1*st*_ and *S*_2*nd*_ are the area of the two consecutive window RFi, and *S_c_* is their colocalized area. A wrapping index of +1 indicates that the RFi in the 1^st^ time window completely wraps around those in the 2^nd^ window, whereas a wrapping index of −1 indicates the reverse (**Fig. S4c**). From our STORM images, we found a WI of −0.88 ± 0.03 in early S-phase, −0.03 ± 0.29 in mid S-phase, and +0.83 ± 0.05 in late S-phase (**Fig. 4f**), clearly illustrating the reversal in the radial directionality of RFi propagation. A similar reversal was observed when the order of the two dye-labeled nucleotides used was swapped (**Fig. S4d**), thereby eliminating the possibility that such propagation pattern could be the consequence of labeling and detection artifacts associated with specific dyes. Moreover, when cells were treated with 5 mM caffeine, a well-known inhibitor of cell cycle checkpoint and bring the origin firing from late to early S-phase^49^, the replication pattern in early S-phase became similar to that in mid S-phase, especially around nuclear periphery and nucleoli (**Fig. 4g**). In addition, the diameter of RFi decreased drastically 338.5±25.6 nm to 137.9±12.6 nm (**Fig. 4h**), and the radial propagation pattern observed for early S-phase was abrogated, as indicated by a near-zero WI with caffeine treatment (**Fig. 4i**).

### CTCF regulates RFi morphology and propagation dynamics

In order to understand the mechanistic basis underlying the unique dynamics of RFi propagation, we remind ourselves of the prominent role that CTCF plays in regulating 3D genome structure through forming chromatin loops, and hypothesize that CTCF-mediated loop structures serve both as structural basis for the replicating DNA as well as spatial constraints for their propagation. To that end, we quantitatively characterized the morphology and propagation dynamics of early S-phase RFi in cells treated with either nonspecific control (NC) siRNA or CTCF siRNA (**Fig. 5a**), and confirmed the downregulation of CTCF both at the levels of mRNA and protein using qPCR, immunostaining, and western blotting, respectively (**Fig. 5a** inset, **Fig. S5a and b**). Downregulation of CTCF enlarged RFi size without significantly reducing their density (**Fig. 5b** and **c**). However, the local density (defined as the number of single-molecule localizations in STORM images that constitute each focus) in the RFi increased significantly (**Fig. 5d**). More importantly, the radially specific propagation pattern of RFi described in **Fig. 4** was drastically diminished by CTCF knockdown (**Fig. 5e**), as evidenced by RFi diameter (**Fig. 5f**) and the extent colocalization ratio between RFi activated during two adjacent labeling windows (**Fig. 5g**), as well as the Wrapping Index (**Fig. 5h**) from −0.79±0.06 to +0.22±0.52. These results strongly suggest that CTCF and the loop structures it mediates act as a key factor underlying RFi morphological dynamics of propagation.

**Figure 5.**
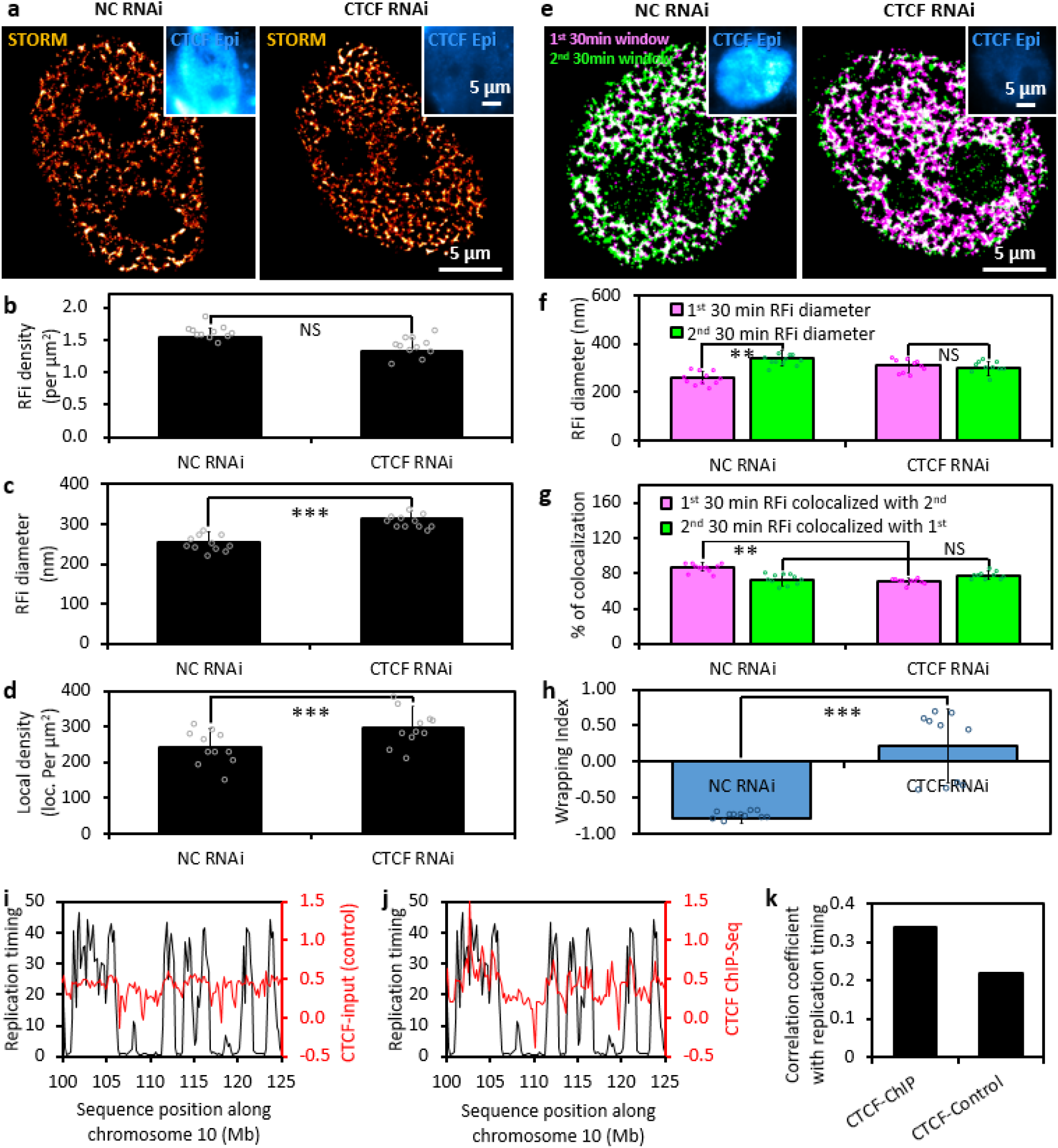
CTCF regulates RFi morphology and spatio-temporal dynamics. **a**, STORM images of RFi labeled at early S-phase transfected with either nonspecific control (NC) siRNA (left) or CTCF siRNA (right). Inset in each image shows the immune-fluorescently labeled for CTCF (blue). See also **Fig. S6** for western blot and qPCR quantification. **b-d**, Density **(b)**, diameter **(c)**, and single-molecule localization density (d) of RFi in cells treated with either NC or CTCF siRNA. **e**, Dual-color STORM images of newly replicated DNA labeled in two consecutive 30 min windows (purple and green, respectively) at the beginning of early S-phase in cells treated with either NC (left) or CTCF siRNA (right). **f**, Diameters of RFi in the first (purple) and second (green) labeling windows. **g**, Fractions of RFi in the first window that colocalized with those in the second window (pink), and vice versa (green). **h**, Wrapping Index between the RFi produced in the two labeling windows. **i, j**, Replication timing profile (black line) along human chromosome 10 mapped onto ChIP-Seq data of either control (**i**,red line) or CTCF (**j**, red line). **k**, Correlation coefficient between replication timing and ChIP-Seq data for both control and CTCF, at the bin size of 200kb. Error bars denote mean ± s.d. p values are determined by Student’s t test; **: p < 0.01; ***: p < 0.001; NS: not significant. (n = 11 cells for **(b)** to **(d)** and **(f)** to **(h)**). Source data are provided as a Source Data file.

To further correlate replication timing, RFi propagation dynamics and CTCF-mediated chromatin structures, we mapped the replication timing profile and ChIP-Seq data along human chromosomes for both control (input) and CTCF (**Fig. 5i** and **j**, showing only a specific region along chromosome 10). The CTCF binding sequences exhibit a high level of correlation with that of the replication timing profile (**Fig. 5k**). These bioinformatic analyses, along with our STORM imaging results, strongly suggest that CTCF-organized chromatin structures play an important role in facilitating the non-random activation of DNA replication and its radial propagation within individual RFi.

### The “CoREP” model for non-random replication activation within RD

To account for the observed link between RFi propagation pattern and the CTCF-mediated chromatin organization, we propose a spatio-temporal model, termed ‘CTCF-organized REplication Propagation’ (CoREP), for the non-random activation and propagation of DNA replication at the sub-RD level (**Fig. 6**). Our model is based on previous findings that a RD contains one or several TADs (each with a mean size of 880 kb^17^), and that each TAD contains 4~6 CTCF-organized DNA loops (each with a mean size of 185 kb^40^). These loops, occupied by promotors, enhancers and active genes, are organized at their anchor sites by CTCF and cohesin (**Fig. 6a**). Our observed spatio-temporal patterns of RFi propagation suggest that replication activation events may take place preferentially at CTCF-mediated loop anchors within each RD, which then propagate to the periphery of the domain (**Fig. 6b**, model I CoREP). Alternatively, it is also possible that DNA replication could be activated at the periphery of each RD, and chromatin structural reorganization then drives the DNA that are replicated earlier to move towards the interior of the domain and those replicated later to the exterior (**Fig. S6**). Whichever is the case, our findings argue against the random activation of replication within RD (**Fig. 6c** and **d**, models II and III). To further validate our model, we simulated the propagation of RFi for each of these three models and calculated the radius of gyration of the simulated RFi (see Methods, Supplementary Materials and **Figs. S7** to **S9** for details). We found that the experimentally observed RFi size (**Fig. 4d**) agree much better with the simulated radius of gyration using the “CoREP” model (**Fig. 6e**) as compared to those using the random firing models based on either looped or random chromatin structures (**Fig. 6f**). These data support a chromatin structure-mediated mechanism of replication activation and propagation inside individual RFi.

**Figure 6.**
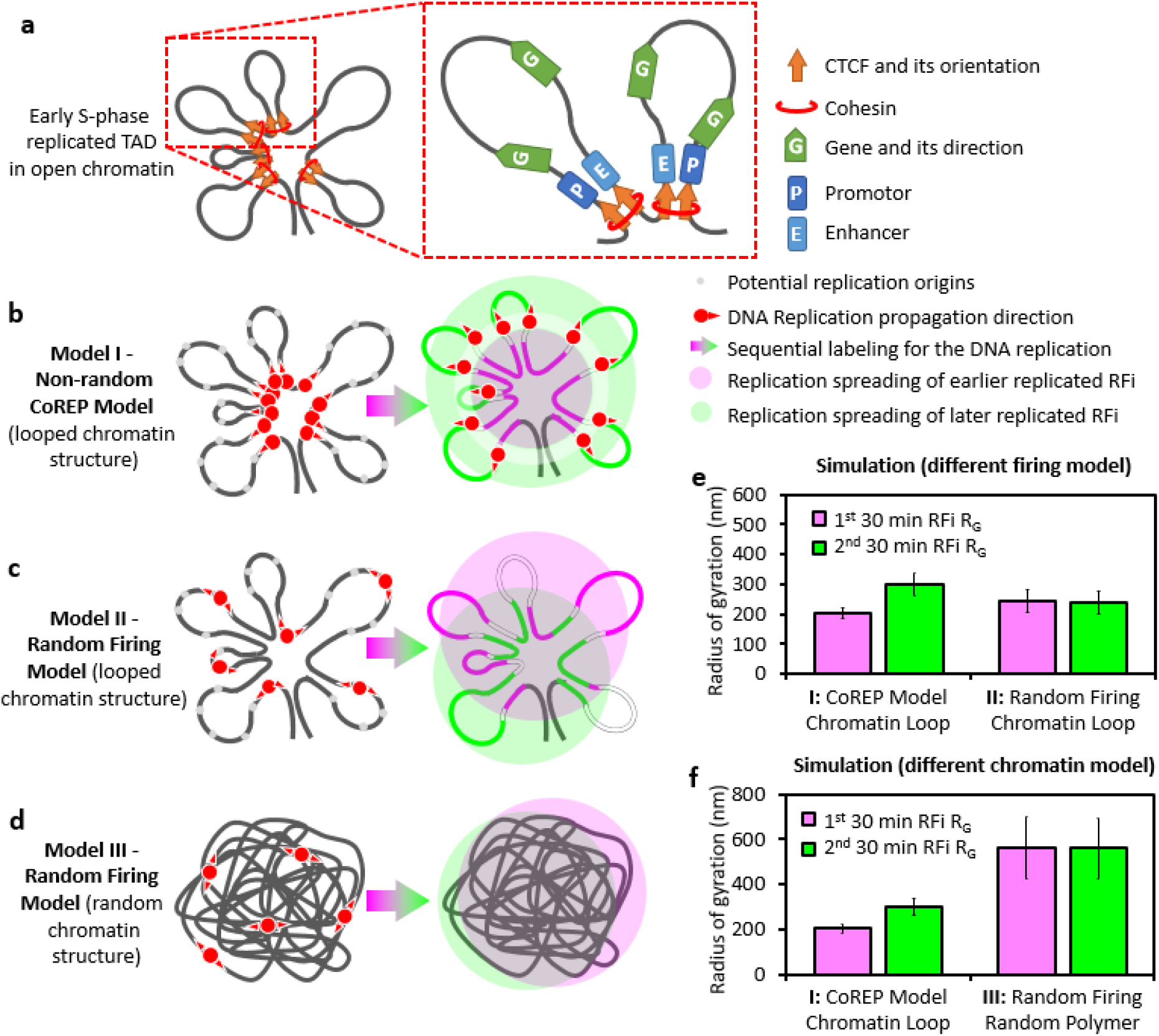
The ‘CTCF-organized REplication Propagation’ (CoREP) model for DNA replication activation and propagation. **a**, Schematic diagram of an early S-phase TAD, organized into loop structures by CTCF and located at the base; each loop contains promotors, enhancers and the active genes. **b-d**, Three models of spatio-temporal replication propagation. **(b)** The CoREP model, in which replication is preferentially activated from inside of CTCF-organized chromatin structure; **(c)** Random firing model with looped chromatin structure; **(d)** Random firing model with random chromatin structure. **e**, Simulated radii of gyration of RFi produced in two consecutive 30-min labeling windows using I: CoREP or II: random firing models. **f**, Comparison of simulated radii of gyration of RFi produced in two consecutive 30-min labeling window using random firing model based on either I: looped or III: random chromatin structures. Error bars denote mean ± s.d. (n = 500 chromatin configurations for **(e)** and **(f)**).

## DISCUSSION

DNA replication in mammalian cells takes place at thousands of foci across the genome, and must be coordinated precisely to ensure the complete and faithful duplication of the genomic information of the cell. One of the key unanswered questions regarding the mechanism of DNA replication concerns how the process is activated, propagated, and regulated in both space and time. While replication activation has been found to correlate with a combination of factors, including G-quadruplex motifs^14^, chromatin modifications^10,50,51^, transcription activity^9^ and more recently early replication control elements (ERCEs)^52^, but no general rule has been established. Previous studies have also suggested that the initiation of replication is regulated at the RD level, but within a RD the choice of replication origins to fire is often believed to be flexible and stochastic^13,14^. Here, we fill a critical gap in our understanding of the spatio-temporal coordination of replication by performing, to our knowledge, the first quantitative characterization of the *in situ* epigenetic environment of RFi inside mammalian cell nucleus across different stages of S-phase. Our super-resolution imaging data reveal a distinct radial propagation pattern of DNA replication within RD, and strongly suggest a non-random selection mechanism for replication activation. The “CoREP” model we proposed further establishes the requirement of CTCF for this process, and points to the essential role of local chromatin environment and structures in DNA replication activation. In that respect, our findings are in line with previous reports of the involvement of CTCF and cohesion in organizing the genome into numerous ‘contact domains’^40,41,53,54^, as well as the enrichment of CTCF and cohesin at TAD boundaries^18^ and replication sites^53^ respectively.

Overall, our work provides a novel spatio-temporal model for replication propagation, and establishes the fundamental framework for further quantitative investigations into the organization and dynamics of DNA replication at the super-resolution level. For example, the relationship between the RFi observed in Cartesian space and RDs in sequence space revealed by replication timing profiles could be probed by co-labeling specific RD sequences using DNA fluorescence *in situ* hybridization (FISH) and measuring their colocalization with RFi. The occurrence of chromatin reorganization or relocation can also be checked by sequentially labeling the same sequence at both G1 and S-phase, while sequential labeling RFi at the same stage in contiguous cell cycles could reveal the extent to which replication originates from the same sites in different cell cycles. Moreover, a recent study using ChIA-PET has revealed that the chromatin structures organized by CTCF/cohesin spatially coordinate RNA polymerase II transcription^41^, pointing to an intriguing direction towards understanding the coupling between replication and transcription. Open chromatin around transcriptionally active promoters could facilitate the selection of replication origins in the vicinity, which might in turn explain previous observations that transcriptionally active genes tend to replicate early while inactive ones tend to replicate late during S-phase^41^. In fact, replication initiation has been found to often occur adjacent to transcription initiation events, and replicator elements can contribute to transcriptional activation^14,55^. By combining the approaches used here with those developed previously for probing RNA polymerase II-mediated transcription^56,57^ and transcription factor binding^58^, we can probe the extent to which replication and transcription are colocalized/segregated in space^59^, thereby generating new insights into the cooperativity between these two critically important nuclear processes.

## METHODS

### Cell culture and synchronization

The HeLa-S3 immortalized cell line (PubMed ID: 5733811) was obtained from Dr. Wei Guo, Department of Biology, University of Pennsylvania. HeLa S3 cells were routinely cultured in high-glucose DMEM (Invitrogen) supplemented with 10% (v/v) fetal bovine serum (Gibco), 100 U/ml penicillin (Invitrogen) and 100 mg/ml streptomycin (Invitrogen) at 37 °C with 5% CO2. The duration of the cell cycle was determined by counting the number of cells every 8 hr for a continuous period of 72 hr. The cells were synchronized to the G1/S boundary with sequential treatment of 2 mM thymidine for 15 hr (first DNA replication block period), normal culture medium for 10 hr (interim release period), and 2 μg/ml Aphidicolin for 15 hr (second DNA replication block period). The synchronization efficiency was determined by measuring the cellular DNA content profile in a population of DAPI-stained cells, fixed at 30 min after release into S-phase, using a fluorescence-activated cell sorter (MoFlo, Beckman Coulter) at excitation wavelength of 340-380 nm.

### RNA interference

HeLa S3 cells were transfected with the following siRNA oligos (GenePharma) using Lipofectamine 2000 (Invitrogen) according to the manufacturer’s instructions: non-specific: CGUACGCGGAAUCUUCGATT (sense) and UCGAAGUAUUCCGCGUACGTT (anti-sense); CTCF: GGAGCCUGCCGUAGAAAUUTT (sense) and AAUUUCUACGGCAGGUCCTC (anti-sense). Cells were harvested 48 hrs after transfection, followed by Western blot and qPCR detection of CTCF.

### Replication foci (RFi) labeling and chromatin marker immuno-staining

At specific times after releasing the cells from G1/S boundary, replication foci in the nucleus were labeled by introducing different pulses (30 min for **Fig.1, 3, 4 and 5**; 10, 30 or 60 min for **Fig. 2**) of thymidine analogues. Dye-labeled dUTPs (either Alexa 647 (Invitrogen), Cy5 (GE Healthcare) or Atto 550 (MolBi Tech)) were delivered into the nucleus via transfection using the FuGENE 6 reagent (Promega), while EdU (Life Technologies), being uncharged, was directly added into the culture medium and subsequently conjugated with a dye using the Click-iT EdU Imaging Kit (Life Technologies) according to the manufacturer’s instructions. When performing two-color labeling, the cells were washed with medium for three times after incubating with the first dye before adding in the second dye. For caffeine treatment, a final concentration of 5 mM caffeine (Sigma-Aldrich) was added into the culture medium during the labeling time.

Upon labeling, the sample was incubated with extraction buffer (0.1 M PIPES pH 7.0, 1 mM EGTA, 1 mM MgCl2 and 0.2% (v/v) Triton X-100 (all from Sigma-Aldrich)) for 60~70 seconds and fixed with 4% (w/v) paraformaldehyde (PFA, Electron Microscopy Science) and 0.1% (v/v) glutaraldehyde (GA, Electron Microscopy Science) in PBS (Sigma-Aldrich) at room temperature for 15 min. To stain the proteins of interest, the cells were blocked and permeabilized with blocking buffer (5% (w/v) BSA (Jackson Immuno Research) and 0.5% (v/v) TritonX-100 (Sigma-Aldrich) in PBS) for 30 min, and incubated with 1~10 μg/ml (according to the manufacturer manual) primary antibodies towards proteins of interest (PCNA (Santa Cruz, mouse monoclonal, sc-56), H3K27ac (Millipore, rabbit polyclonal, 07-360), H3K4me3 (Active Motif, rabbit polyclonal, 39159), CTCF (Abcam, rabbit monoclonal, ab128873), SUZ12 (Bethyl Laboratories, rabbit polyclonal, a302-407a), LaminA/C (Abcam, mouse monoclonal, ab40567), Histone H2B (Santa Cruz, goat polyclonal, sc8650)) in the blocking buffer for 60 min at room temperature (the incubation condition of Histone H2B antibody is 20μg/ml for 10 hrs at 4 °C). After washing with PBS for 3 times, the primary antibody was visualized using 2~5 μg/ml secondary antibody towards specific species of primary antibody labeled with Cy3B (GE Healthcare) or Atto 488 (Sigma-Aldrich) in the blocking buffer for 40 min at room temperature. The sample was post-fixed with 4% (w/v) PFA in PBS for 10 min.

### DNA combing

After labeling with EdU or Dye-dUTP, HeLa cells were harvested and resuspended in PBS to a concentration of 5×10^5^ cells/mL and diluted 4-fold with unlabeled cells at the same concentration. Cells were lysed with lysis buffer (200 mM Tris-HCl at pH 7.5, 50 mM EDTA and 0.5% (w/v) SDS) and added onto a coverslip and let dry for 3~5 min. The coverslip was then tilted up at 15° to allow the DNA to flow down slowly along the coverslip. The coverslip were air-dried and fixed in 3:1 methanol/acetic acid, followed by incubation at 4 ° C overnight.

### Super-resolution imaging and data analysis

STORM imaging was similarly performed as previously described^47,60^. The super-resolution images of Alexa 647-, Cy3B-, Atto 550 and Atto 488-labeled replication domains or proteins of interest were continuously acquired for up to 100,000 frames under the excitation of 647nm, 561nm and 488nm laser at the power density of 3~5 kW/cm^2^ and under the activation of 405nm laser at the power density of 0.5 kW/cm^2^. Bright-field images of 3 μm sized glass beads (Weike Co., Wuhan) placed in the sample dish prior to imaging were acquired 100 ms every 1000 ms for correcting lateral drift between the frames. STORM image analysis, drift correction, RFi quantification, image rendering and images presentation were performed using Insight3 (gift of Prof. Bo Huang from UCSF^60^), custom-written Matlab (2011a, MathWorks) codes, SR-Tesseler (IINS, Interdisciplinary Institute for Neuroscience^32^), and Image J (Image Processing and Analysis in Java). Colocalization was defined as the centroid positions of two structures being not more than 100 nm (3 fold of STORM resolution in cell nucleus) apart from each other. Wrapping Index (WI) was defined as 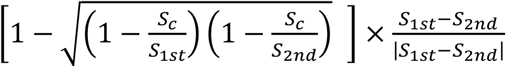, where *S*_1*st*_ and *S*_2*nd*_ are the area of the two consecutive window RFi, and *S_c_* is their colocalized area.

### Simulation of replication foci propagation

To understand the intriguing spatio-temporal propagation pattern of DNA replication that was sequentially labeled (**Fig. 4** and **5**), we simulated the spatial organization of chromatin at the molecular level using sets of chromatin chains with internal loops in the 3D space. The theoretical modeling includes: (1) Sampling of short chromatin chains using the coarse-grained nucleosome model and DNA model that have similar geometry and flexibility compared to the real system. (2) Connections of short chromatin chains into long chains with internal loops to simulate the chromatin structures organized by CTCF/cohesin. (3) Split of a long chromatin chain into two segments to represent DNA replicated during the two consecutive labeling time windows in **Figure 5**. See supplementary materials for details.

### Sequencing data and correlation analyses

We compared replication timing and CTCF ChIP-seq data of Hela-S3 cells to show their correlation. For replication timing data, we used the percentage-normalized signal profile from ENCODE downloads of UCSC (http://genome.ucsc.edu/ENCODE/) where replication signal at 1 kb intervals is a percentage of normalized +/-25 kb tag densities. CTCF ChIP-seq data is also from ENCODE downloads of UCSC but was remapped to hg19 using bowtie and reanalyzed using MACS. All chromosomes were divided into bins of different lengths (1k, 5k, 10k, 50k, 100k, 200k, 500k, 1M, 2M, 5M) at the same position. For both replication timing and CTCF ChIP-seq profiles, we calculated median of each bin. And then for each length, correlation coefficient was calculated using Pearson method in R.

### Code availability

The code for the polymer simulation can be downloaded at http://wikisend.com/download/164174/code.tar

### Data Availability Statement

The datasets generated during and/or analysed during the current study are available from the corresponding author on reasonable request.

## AUTHOR CONTRIBUTIONS

Y.S. and X.S.X. conceived the project; Y.S., Q.P.S. and Z.W.Z. designed the experiments; Q.P.S., Z.W.Z., M.D., W.Z., Y.L., and R.L. performed the wet lab experiments; Q.P.S performed all the cell labeling and STORM imaging experiments; Q.P.S. and Y.S. wrote the Matlab codes; Q.P.S. and Z.W.Z. performed data analysis; L.M. and Y.Q.G. performed the simulation analysis; M.L. performed the bioinformatics analysis; Y.S., X.S.X. and Y.G. supervised the project; Q.P.S., Z.W.Z., Y.G., X.S.X. and Y.S. wrote the manuscript.

## ACKNOWLEDGEMENTS

We thank Prof. Xiaowei Zhuang and Dr. Hazen Babcock (Harvard University) for help with optical setup of the STORM microscope and Prof. Bo Huang (University of California, San Francisco) for Insight3 software. We thank the Core Facilities at School of Life Sciences, Peking University for imaging support. This work is supported by grants National Science Scholarship from the Agency for Science, Technology and Research (A*STAR) of Singapore for Z.W.Z, the National Science Foundation of China 21573013, 21390412, 31271423, and 31327901, 863 Program SS2015AA020406 and CAS Interdisciplinary Innovation Team for Y.S.

## REFERENCES

1 Perumal, S. K., Yue, H., Hu, Z., Spiering, M. M. & Benkovic, S. J. Single-molecule studies of DNA replisome function. Biochim. Biophys. Acta 1804, 1094–1112 (2010).

2 van Oijen, A. M. & Loparo, J. J. Single-molecule studies of the replisome. Annual review of biophysics 39, 429–448, doi:10.1146/annurev.biophys.093008.131327 (2010).

3 Jacob, F. The replicon: Thirty years later. Cold Spring Harb. Symp. Quant. Biol. 58, 383–387 (1993).

4 Leonard, A. C. & Mechali, M. DNA replication origins. Cold Spring Harbor perspectives in biology 5, a010116, doi:10.1101/cshperspect.a010116 (2013).

5 Hozak, P., Hassan, A. B., Jackson, D. A. & Cook, P. R. Visualization of replication factories attached to nucleoskeleton. Cell 73, 361–373 (1993).

6 Rhind, N. & Gilbert, D. M. DNA replication timing. Cold Spring Harbor perspectives in biology 5, a010132, doi:10.1101/cshperspect.a010132 (2013).

7 Hiratani, I. et al. Global reorganization of replication domains during embryonic stem cell differentiation. PLoS biology 6, e245, doi:10.1371/journal.pbio.0060245 (2008).

8 Ryba, T. et al. Evolutionarily conserved replication timing profiles predict long-range chromatin interactions and distinguish closely related cell types. Genome research 20, 761–770, doi:10.1101/gr.099655.109 (2010).

9 Dimitrova, D. S. & Gilbert, D. M. The spatial position and replication timing of chromosomal domains are both established in early G1 phase. Molecular cell 4, 983–993 (1999).

10 Fragkos, M., Ganier, O., Coulombe, P. & Mechali, M. DNA replication origin activation in space and time. Nature reviews. Molecular cell biology 16, 360–374, doi:10.1038/nrm4002 (2015).

11 Besnard, E. et al. Unraveling cell type-specific and reprogrammable human replication origin signatures associated with G-quadruplex consensus motifs. Nature structural & molecular biology 19, 837–844, doi:10.1038/nsmb.2339 (2012).

12 Ferguson, B. M. & Fangman, W. L. A position effect on the time of replication origin activation in yeast. Cell 68, 333–339 (1992).

13 Rivera-Mulia, J. C. & Gilbert, D. M. Replicating Large Genomes: Divide and Conquer. Molecular cell 62, 756–765, doi:10.1016/j.molcel.2016.05.007 (2016).

14 Cayrou, C. et al. Genome-scale analysis of metazoan replication origins reveals their organization in specific but flexible sites defined by conserved features. Genome research 21, 1438–1449, doi:10.1101/gr.121830.111 (2011).

15 Kaykov, A. & Nurse, P. The spatial and temporal organization of origin firing during the S-phase of fission yeast. Genome research 25, 391–401, doi:10.1101/gr.180372.114 (2015).

16 Jackson, D. A. & Pombo, A. Replicon clusters are stable units of chromosome structure: evidence that nuclear organization contributes to the efficient activation and propagation of S phase in human cells. The Journal of cell biology 140, 1285–1295 (1998).

17 Pope, B. D. et al. Topologically associating domains are stable units of replication-timing regulation. Nature 515, 402–405, doi:10.1038/nature13986 (2014).

18 Dixon, J. R. et al. Topological domains in mammalian genomes identified by analysis of chromatin interactions. Nature 485, 376–380, doi:10.1038/nature11082 (2012).

19 Ricci, M. A., Manzo, C., Garcia-Parajo, M. F., Lakadamyali, M. & Cosma, M. P. Chromatin fibers are formed by heterogeneous groups of nucleosomes in vivo. Cell 160, 1145–1158, doi:10.1016/j.cell.2015.01.054 (2015).

20 Boettiger, A. N. et al. Super-resolution imaging reveals distinct chromatin folding for different epigenetic states. Nature 529, 418–422, doi:10.1038/nature16496 (2016).

21 Xiang, W. et al. Correlative live and super-resolution imaging reveals the dynamic structure of replication domains. The Journal of cell biology 217, 1973–1984, doi:10.1083/jcb.201709074 (2018).

22 Xu, J. et al. Super-Resolution Imaging of Higher-Order Chromatin Structures at Different Epigenomic States in Single Mammalian Cells. Cell reports 24, 873–882, doi:10.1016/j.celrep.2018.06.085 (2018).

23 Jackson, D. A. S-Phase progression in synchronized human cells. Exp. Cell Res. 220, 62–70 (1995).

24 Ma, H. T. & Poon, R. Y. C. Synchronization of HeLa cells. Methods Mol. Bio. 761, 151–161, doi:10.1007/978-1-61779-182-6_10 (2011).

25 Dimitrova, D. S. Visualization of DNA replication sites in mammalian nuclei. Methods Mol. Bio. 521, 413–436, doi:10.1007/978-1-60327-815-7_23 (2009).

26 Koberna, K. et al. Electron microscopy of DNA replication in 3-D: evidence for similar-sized replication foci throughout S-phase. Journal of cellular biochemistry 94, 126–138, doi:10.1002/jcb.20300 (2005).

27 Ma, H. et al. Spatial and temporal dynamics of DNA replication sites in mammalian cells. The Journal of cell biology 143, 1415–1425 (1998).

28 Baddeley, D. et al. Measurement of replication structures at the nanometer scale using super-resolution light microscopy. Nucleic acids research 38, e8, doi:10.1093/nar/gkp901 (2010).

29 Chagin, V. O. et al. 4D Visualization of replication foci in mammalian cells corresponding to individual replicons. Nature communications 7, 11231, doi:10.1038/ncomms11231 (2016).

30 Cseresnyes, Z., Schwarz, U. & Green, C. M. Analysis of replication factories in human cells by super-resolution light microscopy. BMC cell biology 10, 88, doi:10.1186/1471-2121-10-88 (2009).

31 Triemer, T. et al. Super-resolution imaging of individual replication forks reveals unexpected prodrug resistance mechanism. Proceedings of the National Academy of Sciences of the United States of America 115, E1366–e1373, doi:10.1073/pnas.1714790115 (2018).

32 Levet, F. et al. SR-Tesseler: a method to segment and quantify localization-based super-resolution microscopy data. Nature methods 12, 1065–1071, doi:10.1038/nmeth.3579 (2015).

33 Schwob, E. et al. in DNA Replication: Methods and Protocols (eds Sonya Vengrova & Z. Jacob Dalgaard) 673–687 (Humana Press, 2009).

34 Malinsky, J. et al. The supply of exogenous deoxyribonucleotides accelerates the speed of the replication fork in early S-phase. Journal of cell science 114, 747–750 (2001).

35 Rhind, N. DNA replication timing: random thoughts about origin firing. Nature cell biology 8, 1313–1316, doi:10.1038/ncb1206-1313 (2006).

36 Chagin, V. O., Stear, J. H. & Cardoso, M. C. Organization of DNA replication. Cold Spring Harbor perspectives in biology 2, a000737, doi:10.1101/cshperspect.a000737 (2010).

37 Michalet, X. et al. Dynamic molecular combing: stretching the whole human genome for high-resolution studies. Science 277, 1518–1523 (1997).

38 Wang, S. et al. Spatial organization of chromatin domains and compartments in single chromosomes. Science 353, 598–602, doi:10.1126/science.aaf8084 (2016).

39 Ong, C. T. & Corces, V. G. CTCF: an architectural protein bridging genome topology and function. Nature reviews. Genetics 15, 234–246, doi:10.1038/nrg3663 (2014).

40 Rao, S. S. et al. A 3D map of the human genome at kilobase resolution reveals principles of chromatin looping. Cell 159, 1665–1680, doi:10.1016/j.cell.2014.11.021 (2014).

41 Tang, Z. et al. CTCF-Mediated Human 3D Genome Architecture Reveals Chromatin Topology for Transcription. Cell, doi:10.1016/j.cell.2015.11.024 (2015).

42 Barski, A. et al. High-Resolution Profiling of Histone Methylations in the Human Genome. Cell 129, 823–837, doi:10.1016/j.cell.2007.05.009 (2007).

43 Creyghton, M. P. et al. Histone H3K27ac separates active from poised enhancers and predicts developmental state. Proceedings of the National Academy of Sciences of the United States of America 107, 21931–21936, doi:10.1073/pnas.1016071107 (2010).

44 Peric-Hupkes, D. & van Steensel, B. Role of the nuclear lamina in genome organization and gene expression. Cold Spring Harb. Symp. Quant. Biol. 75, 517–524, doi:10.1101/sqb.2010.75.014 (2010).

45 Cao, R. & Zhang, Y. SUZ12 Is Required for Both the Histone Methyltransferase Activity and the Silencing Function of the EED-EZH2 Complex. Molecular Cell 15, 57–67, doi:10.1016/j.molcel.2004.06.020 (2004).

46 Rinn, J. L. et al. Functional Demarcation of Active and Silent Chromatin Domains in Human HOX Loci by Noncoding RNAs. Cell 129, 1311–1323, doi:10.1016/j.cell.2007.05.022 (2007).

47 Wang, G. et al. PTEN regulates RPA1 and protects DNA replication forks. Cell research 25, 1189–1204, doi:10.1038/cr.2015.115 (2015).

48 Guelen, L. et al. Domain organization of human chromosomes revealed by mapping of nuclear lamina interactions. Nature 453, 948–951, doi:10.1038/nature06947 (2008).

49 Marheineke, K. & Hyrien, O. Control of replication origin density and firing time in Xenopus egg extracts: role of a caffeine-sensitive, ATR-dependent checkpoint. The Journal of biological chemistry 279, 28071–28081, doi:10.1074/jbc.M401574200 (2004).

50 Berbenetz, N. M., Nislow, C. & Brown, G. W. Diversity of Eukaryotic DNA Replication Origins Revealed by Genome-Wide Analysis of Chromatin Structure. PLoS Genet 6, e1001092, doi:10.1371/journal.pgen.1001092 (2010).

51 Eaton, M. L., Galani, K., Kang, S., Bell, S. P. & MacAlpine, D. M. Conserved nucleosome positioning defines replication origins. Genes & Development 24, 748–753, doi:10.1101/gad.1913210 (2010).

52 Sima, J. et al. Identifying cis Elements for Spatiotemporal Control of Mammalian DNA Replication. Cell, doi:10.1016/j.cell.2018.11.036 (2018).

53 Guillou, E. et al. Cohesin organizes chromatin loops at DNA replication factories. Genes & development 24, 2812–2822, doi:10.1101/gad.608210 (2010).

54 Zuin, J. et al. Cohesin and CTCF differentially affect chromatin architecture and gene expression in human cells. Proceedings of the National Academy of Sciences of the United States of America 111, 996–1001, doi:10.1073/pnas.1317788111 (2014).

55 Martin, M. M. et al. Genome-wide depletion of replication initiation events in highly transcribed regions. Genome research 21, 1822–1832, doi:10.1101/gr.124644.111 (2011).

56 Zhao, Z. W. et al. Spatial organization of RNA polymerase II inside a mammalian cell nucleus revealed by reflected light-sheet super-resolution microscopy. Proc. Natl. Acad. Sci. U.S.A. 111, 681–686, doi:10.1073/pnas.1318496111 (2014).

57 Chen, X. et al. Study of RNA Polymerase II Clustering inside Live-Cell Nuclei Using Bayesian Nanoscopy. ACS Nano 10, 2447–2454, doi:10.1021/acsnano.5b07257 (2016).

58 Gebhardt, J. C. et al. Single-molecule imaging of transcription factor binding to DNA in live mammalian cells. Nature methods 10, 421–426, doi:10.1038/nmeth.2411 (2013).

59 Chakalova, L., Debrand, E., Mitchell, J. A., Osborne, C. S. & Fraser, P. Replication and transcription: Shaping the landscape of the genome. Nat Rev Genet 6, 669–677 (2005).

60 Huang, B., Wang, W., Bates, M. & Zhuang, X. Three-dimensional super-resolution imaging by stochastic optical reconstruction microscopy. Science 319, 810–813, doi:10.1126/science.1153529 (2008).

