## Supplementary Materials for "CTCF-mediated Chromatin Structures Dictate the Spatio-temporal Propagation of Replication Foci"

**This PDF file includes:**

**Figs. S1 to S9**

**Supplementary Text**

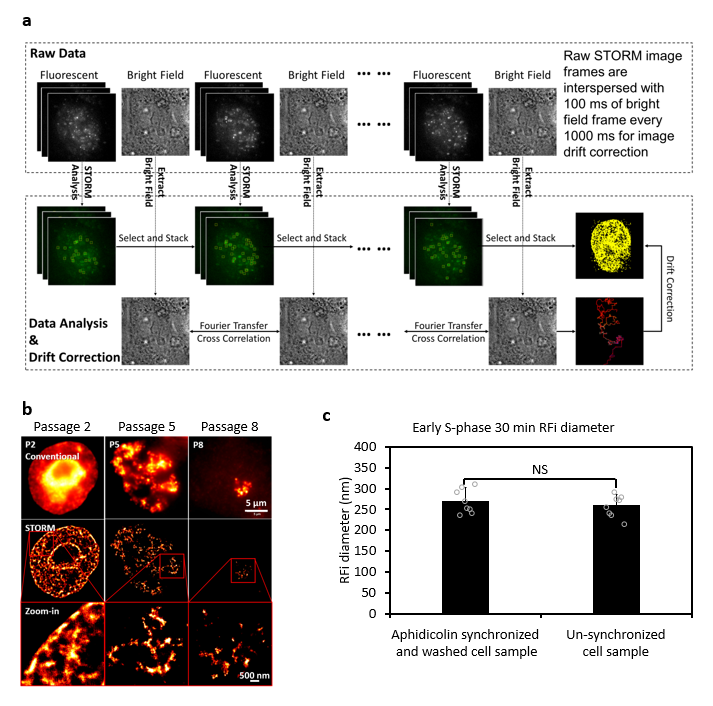

***Figure S1 | Metabolic labeling of replication foci (RFi), STORM imaging procedure and impact of cell cycle synchronization on RFi size. a,*** *Workflow of STORM imaging, drift correction and data analysis for reconstructing super-resolution images.* ***b,*** *Thymidine analogue EdU labeling does not affect cell growth, P2 means passage 2, and so are the P5 and P8. By the 8^th^ passages, there is only one labeled chromosome in the nucleus.* ***c,*** *Early S-phase RFi diameters in Aphidicolin synchronized and released cells (30 min EdU labelling after drug releasing) or non-synchronized cells (30 min EdU labelled without drug treatment, cells were selected that show patterns of typical early S-phase in Fig. 1) indicate that the cell cycle synchronization using Aphidicolin doesn’t affect the RFi diameter. Error bars denote mean ± s.d. p values are determined by Student’s t test; NS: not significant. (n = 8 cells for (****c****)). Source data are provided as a Source Data file.*

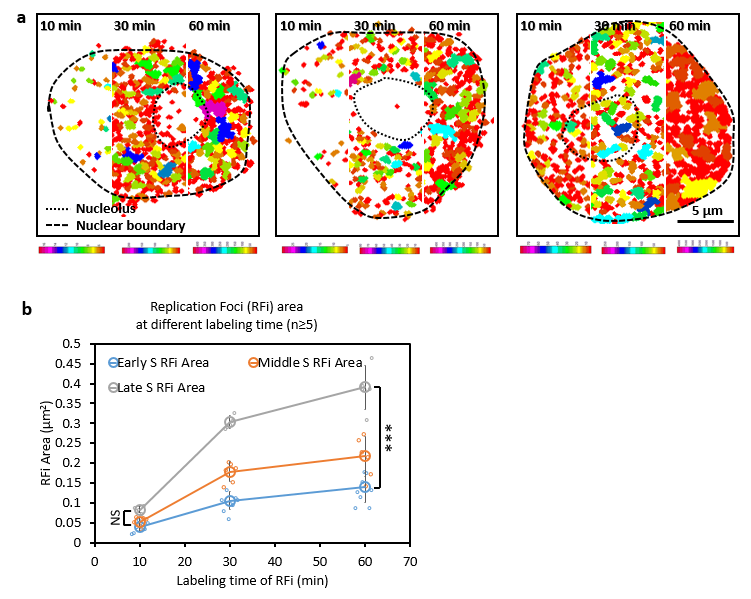

***Figure S2 | Expansion of RFi size informs on variations in replication speed across S-phase. a,*** *RFi of cells with their newly replicated DNA labeled for 10 min, 30 min or 60 min respectively at the beginning of early (left), mid (middle), and late (right) S-phase, related to* ***Figs. 2d*** *to* ***f****. The color coding means the number of dots in a single replication focus.* ***b,*** *Plot showing the mean area of all RFi identified in* ***Figs. 2d to f*** *as a function of labeling duration. Error bars denote mean ± s.d. p values are determined by Student’s t test; NS: not significant, ***: p < 0.001. (n ≥ 5~8 cells for (****a****) and (****b****)). Source data are provided as a Source Data file.*

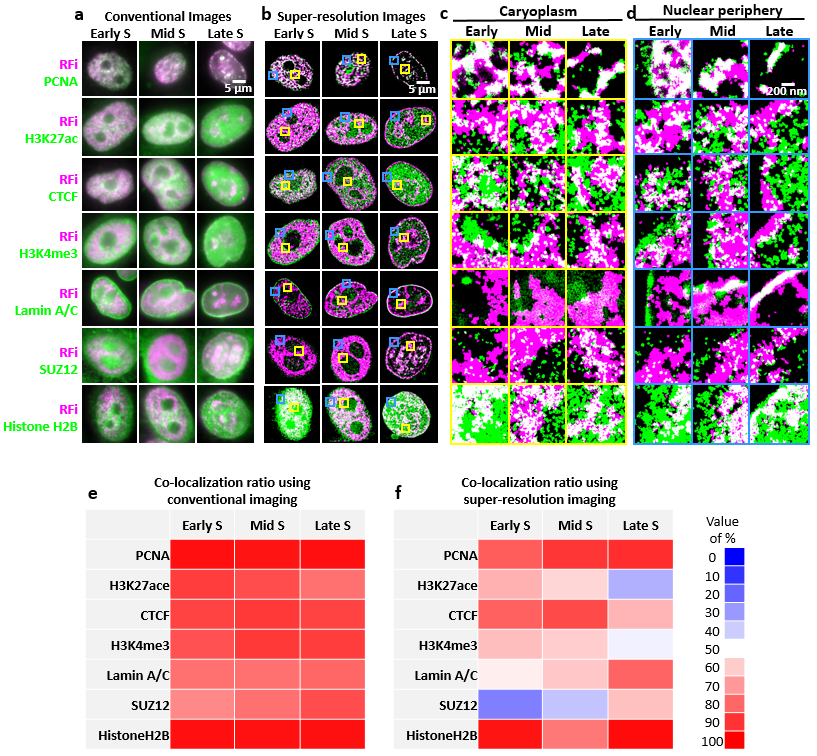

***Figure S3 | Comparison of two-color epi images and zoom-in STORM images of RFi and related nuclear factors and quantification of their colocalization between RFi and related nuclear factors. a,*** *Conventional images of the colocalization between RFi and the seven nuclear factors across different stages of S-phase, related to* ***Fig. 3****.* ***b,*** *STORM images of (A) and inset area for zoom-in.* ***c-d,*** *Caryoplasm (****c****, yellow boxes) and nuclear periphery (****d****, blue boxes) images in (****b****).* ***e, f,*** *Colocalization heat map for conventional (****e****) and STORM images (****f****) of (****a****) and (****b****). Each co-localization ratio is the mean value of 5~8 cells.*

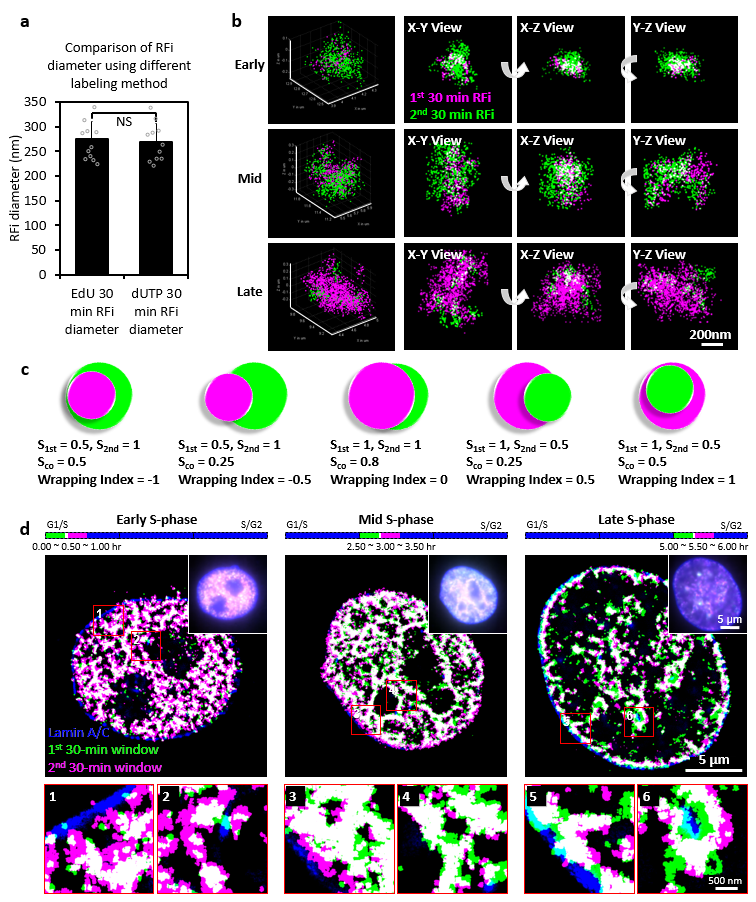

***Figure S4 | Two-color labeling and imaging of RFi reveals a spatio-temporal propagation pattern that reverses directionality from early to late S-phase. a,*** *Comparison of RFi diameter using different nucleotides analogues.* ***b,*** *Localizations constituting a representative RFi from early, mid and late S-phase are displayed in 3D, clearly demonstrating the reversal in the relative spatial spread between the RFi of the two labelling periods, related to* ***Fig. 4****.* ***c,*** *Schematics and definition of wrapping index for RFi in two consecutive labeling time window.* $\left[ 1-\sqrt{\left( 1-\frac{S_{c}}{S_{1st}} \right)\left( 1-\frac{S_{c}}{S_{2nd}} \right)} \right]\times\frac{S_{1st}-S_{2nd}}{\left| S_{1st}-S_{2nd} \right|}$*.* ***d,*** *Newly replicated DNA in two consecutive 30 min periods at the beginning of early, mid and late S-phase were labeled (green and pink), together with lamin A/C (blue) to demarcate the nuclear boundary of each cell. The two dyes used for labeling RFi were added in the opposite order from that used in* ***Fig. 4****. Multi-color STORM images of single labeled nuclei are shown in comparison with epi images of the same nuclei (top right corner), and bars at the top indicate the relative temporal position of the labeling periods during the respective stages of S-phase (not drawn to scale). Insets (numbered 1 through 6) show zoomed-in areas (in red boxes) with enhanced contrast between the two colors, showing the same spreading patterns as in* ***Fig. 4****, thereby eliminating the possibility that such a trend could be the consequence of labeling and detection artifacts associated with specific dyes. Error bars denote mean ± s.d. p values are determined by Student’s t test; NS: not significant. (n = 11 cells for (****a****)). Source data are provided as a Source Data file.*

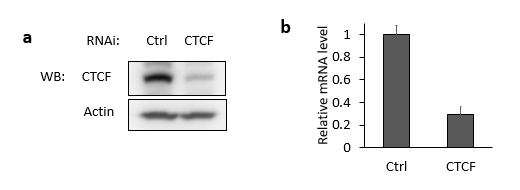

***Figure S5 | Quantification analysis for CTCF RNAi and colocalization of CTCF with replication activation. a, b,*** *Western blot and qPCR towards CTCF are quantified in NC and CTCF RNAi cells, related to* ***Fig. 5****. qPCR was performed in triplicate and the data represent the mean ± s.d. This experiment was repeated 3 times.*

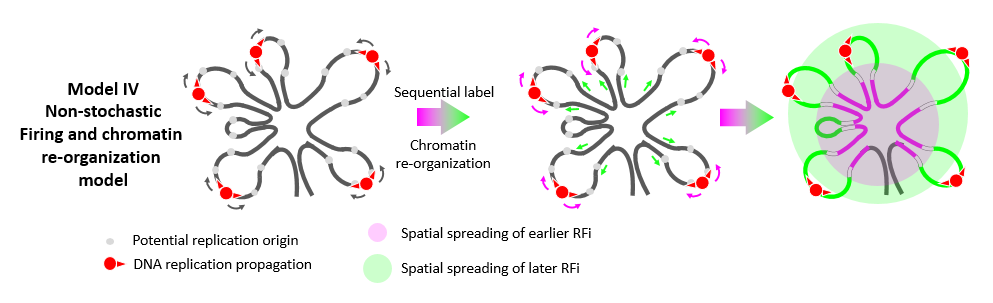

***Figure S6 | Non-stochastic firing and chromatin re-organization model.*** *DNA replication could activate at the surface of RD and chromatin structure reorganization drives DNA that are replicated earlier to move inside and those replicated later to move outside*

**Simulation of replication foci spreading**

To understand the intriguing spatio-temporal propagation pattern of DNA replication that was sequentially labeled (**Fig. 4**), we simulated the spatial organization of chromatin at the molecular level using sets of chromatin chains with internal loops in the 3D space. The theoretical modeling includes: (1) Sampling of short chromatin chains using the coarse-grained nucleosome model and DNA model that have similar geometry and flexibility compared to the real system. (2) Connections of short chromatin chains into long chains with internal loops to simulate the chromatin structures organized by CTCF/cohesin. (3) Split of a long chromatin chain into two segments to represent DNA replicated during the two consecutive labeling time windows in **Fig. 4**.

1. **Sampling of short chromatin chains with coarse-grained model**

In the simulation, we first decomposed a chromatin into nucleosomes and linker DNA chains, which are respectively described by Four-Sphere (FS) and Worm-Like-Chain (WLC) models (shown in **Fig. S7**) and then used these coarse-grained models as building blocks to construct short chromatin chains. In the FS model, one nucleosome is coarse-grained as the coalescence of four identical spheres with a diameter of 5.5 nm. Centers of four spheres are aligned at four vertices of a square with a diagonal length of 5.5 nm. With these settings, the FS model roughly depicts a cylinder-shaped object with a diameter of 11 nm and a thickness of 5.5 nm. Terminal segments of linker DNA chains are placed tangent to the side surface and parallel to the top surface of the cylinder (panel C and D in **Fig. S7**). Two tangent points are located at 1/4 and 3/4 positions along the cylinder axis, respectively. The angle formed by two contact points and the central axis is 120°, roughly consistent with the wrapping of 1.67 turns of DNA in a nucleosome. For the linker DNA, we adopted the WLC model by coarse-graining DNA chains into a sequence of DNA beads with a diameter of 2 nm, in which each bead roughly corresponds to 6 bp in biological system. The DNA beads, sharing the similar diameter with DNA double-helix structure, ensures similar role of steric effect of DNA chains. Potential energy is added to restrict the bending of DNA chains with the form of $E=\sum_{i} \frac{g}{2}\left( \pi-\theta_{i} \right)^{2}$, in which $\theta_{i}$ was the ith bending angle (in the unit of radius) constructed by three neighbor beads along the chain, and g represents the harmonic constant that is to reproduce the flexibility of the chain. In our system, the value of g is set as 60.68 kJ/(mol*rad^2^) to give a DNA persistence length of 50 nm at 300K. The value of the bending angle is set as π at the end of DNA chains to mimic the restriction role of linker histone. For bending angles located in other positions, their values are chosen according to the Boltzmann distribution. In our model, we assumed that each linker DNA chain initially contains 8 DNA beads (or 42 bp) throughout the chromatin and the increase of linker DNA length comes from the exposure of unwrapped DNA if its neighbor nucleosome is removed. Thus, the length of linker DNA can be expressed as 8+33*n* with the probability $P\left( n \right)=\left( 1-\alpha\right)\alpha^{n}$, in which *n* is the number of nucleosomes that are removed in a row and *α* is the removal percentage of nucleosomes in the system.

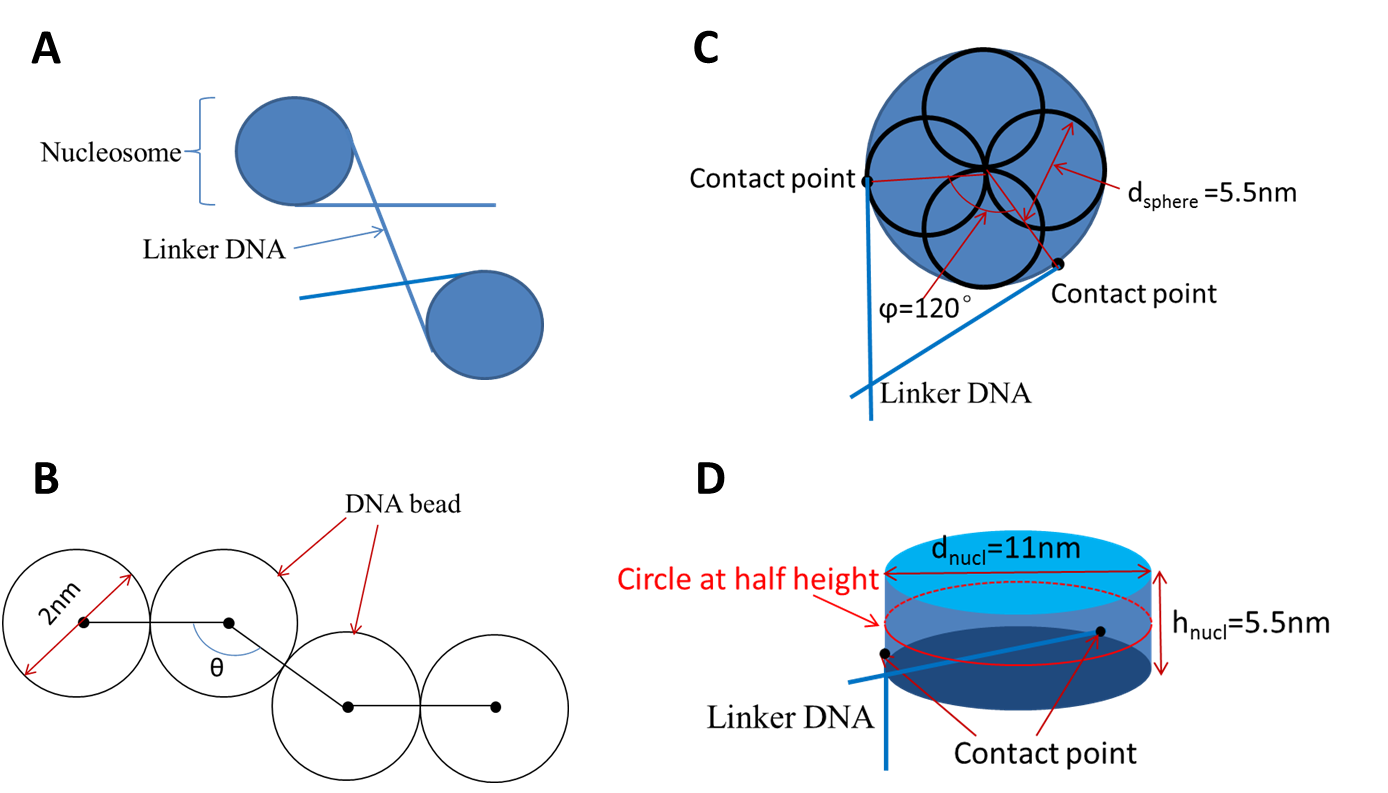

***Figure S7 | Coarse-grained model for chromatin modeling. a,*** *Schematic illustration for the decomposition of chromatin into nucleosomes and a DNA linker that connects them.* ***b,*** *Worm-Like-Chain model for linker DNA.* ***c,*** *Top views of Four-Sphere model. Four spheres in this model are partially overlapped in space. The blue circle represents the projection of the cylinder-shape profile and linker DNA is tangent to the side surface of the cylinder at the contact point.* ***d,*** *Side view of the cylinder profile for Four-Sphere model. Contact points of linker DNA are placed with different height to the top surface.*

Based on the coarse-grained models, the sampling of chromatin chains was conducted by iteratively adding nucleosomes and linker DNAs through a self-avoiding random walk, which was performed in three steps. (1) One nucleosome with two neighbor linker DNA chains is randomly placed in space as the one terminus of chromatin chain. (2) Another nucleosome together with its linker DNA is added to the existing chromatin chain. The new nucleosome is added to the end of the previous linker DNA chain with a random orientation. Overlapping between structural elements is avoided in the addition of new nucleosomes and linkers. (3) The second step is repeated until a desired length of chromatin is reached. In our simulation, 10 sets of short chromatin chains are simulated with nucleosome removal percentage *α* varying from 0.05 to 0.5 with an interval of 0.05. In each set, 1000 chromatin chains of 30 kb long and 1000 chromatin chains of 1 kb long are generated as the basic components for long chromatin chains.

2. **Connecting short chromatin chains into long chromatin chains with internal loops**

To generate target chromatin chains with internal loops, the short chromatin chains are firstly combined to form loop-like chromatins which are further connected to construct long chromatin chains (shown in **Fig. S8**). In the first step, six 30 kb chromatin chains are randomly picked out from 1000 samples and are connected to form one 180 kb chromatin by linker DNA with length of 41 DNA beads (the same length of linker DNA with one nucleosome removed). The 180 kb chromatin resembles the loop-like structure with an end-to-end distance less than 50 nm. A total of 200 chromatin chains of 180 kb are generated in the first step. In the second step, five randomly selected 180 kb chromatins are further connected together by four 1 kb chromatin chains to construct a 900 kb chromatin. Random walk with self-avoidance was used in these processes. After the linkage of short chromatin chains, 500 chromatin chains of 900kb were generated for each chromatin set.

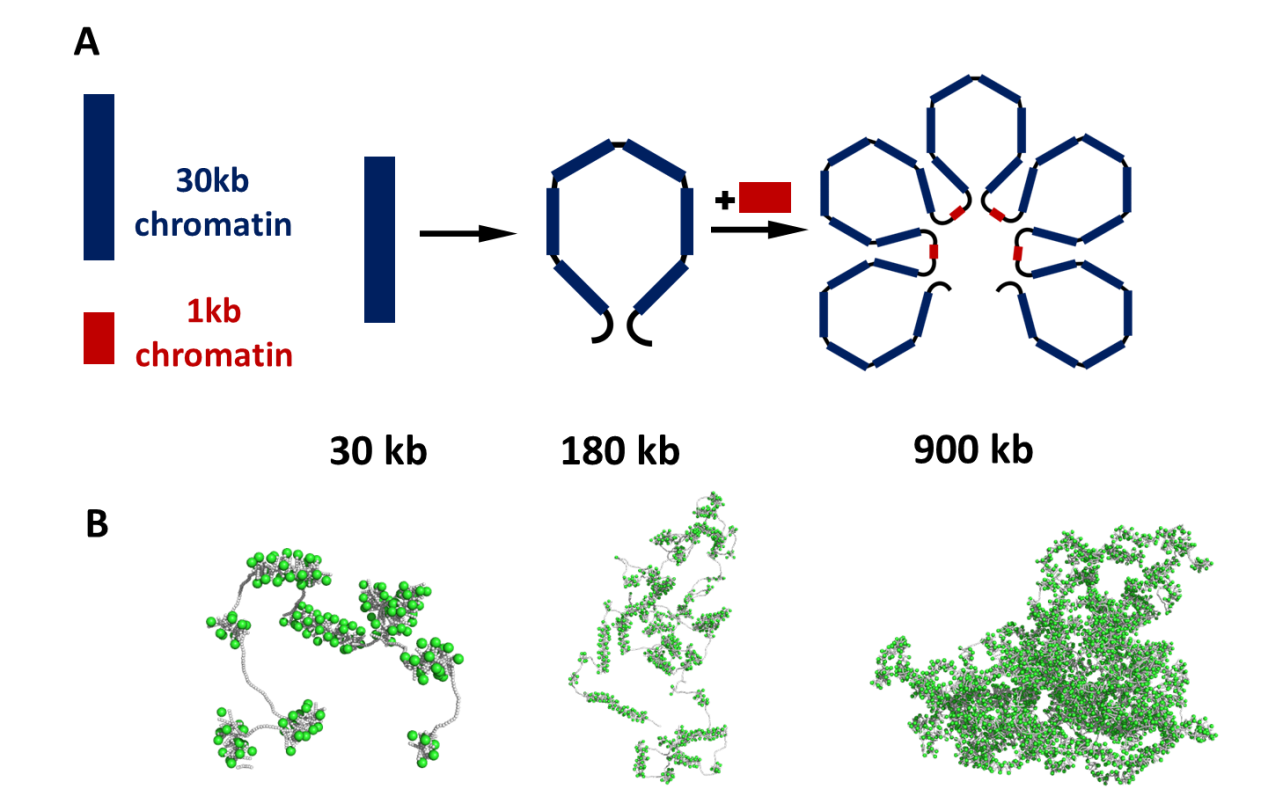

***Figure S8 | Combination of short chromatin chains into 900 kb chromatin chains with internal loops.*** ***a,*** *Schematic flow of the combination process.* ***b,*** *Representative conformation of chromatin chains with length of 30 kb (left), 180 kb (middle) and 900 kb (right). In this plot, nucleosomes are shown in green spheres and DNA beads are shown in white spheres.*

3. **Analysis on the spatial distribution of simulated chromatin**

Starting from different sets of chromatin chains with internal loops, we then try to simulate the spatial organization of chromatins that are replicated during the two consecutive labeling time windows in **Fig. 4**.

In our simulation, we propose two possible models in the replication process. In the first case, we propose the “CoREP” model, in which non-random, selective activation of replication origins occur at the loop-anchors. In contrast, the second model assumes a random activation of replication origins. For each model, we calculated the radius of gyration for the chromatin DNA replicated in the two consecutive labeling time windows (**Fig. S9**). The results show that in the first model, the radius of gyration of chromatin in the later replication time window is significantly larger than that of chromatin in the earlier replication time window. While in the second model, the radii of gyration of chromatin segments are similar in both time windows. Clearly, simulation based on the first model is more consistent with the experimental observations, supporting a non-stochastic, selective mechanism of replication origin activation.

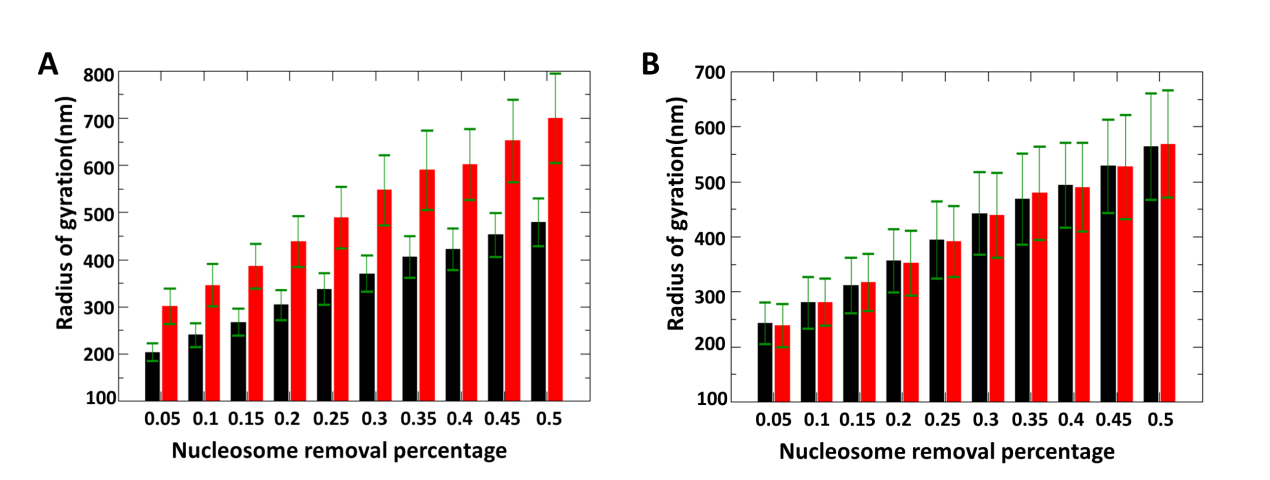

***Figure S9 | Comparison of radii of gyration for chromatin segments replicated in two consecutive replication time windows, simulated by selective (a) and random (b) activation models, respectively. The linker DNA length is adjusted by setting different nucleosome removal percentage α.*** *Black bars represent the radii of gyration in the first replication labeling time window and red bars represent those in the second replication labeling time window. Error bars stand for the standard deviation for the distribution of radius values. Error bars denote mean ± s.d. (n = 500 chromatin configurations)****.*** *Source data are provided as a Source Data file.*
